# Self-regulation of PIN1-driven auxin transport by cell surface-based auxin signaling in *Arabidopsis*

**DOI:** 10.1101/2022.11.30.518523

**Authors:** Jiacheng Wang, Mingzeng Chang, Rongfeng Huang, Michelle Gallei, Jiřn Friml, Yongqiang Yu, Mingzhang Wen, Zhenbiao Yang, Tongda Xu

## Abstract

PIN-FORMED (PIN)-driven auxin transport contributes to establishing and maintaining a dynamic auxin concentration gradient alongside plant tissues, which drives the majority of developmental processes in plants. To maintain developmental plasticity in ever-changing environments, plants have evolved self-organizing feedback machinery between auxin signaling and its transport, which has been shown to play essential roles in many fundamental plant processes. However, the molecular mechanism behind this mutual regulation has not yet been clarified. Here, we identified a cell surface-triggered auxin signaling that regulates the PIN1-mediated auxin efflux and further developmental patterning in *Arabidopsis*. Auxin was able to stimulate PIN1 phosphorylation in plants through transmembrane kinases (TMKs), key components in auxin signaling, at the plasma membrane. TMK1 and TMK4 directly interacted with and phosphorylated PIN1 and functioned redundantly in the regulation of PIN1 polarity in plant cells. The phosphorylation sites in PIN1 proteins, targeted by both auxin and TMKs, were required for PIN1 trafficking and polarity, which further controlled auxin responses and downstream developmental patterning in *Arabidopsis*. Therefore, our findings provide a direct mechanism for the self-regulation between auxin signaling and transport that drives the auxin flows and proper development in plants.

## Main

Auxin regulates virtually every aspect of plant development through its asymmetric distribution, which is based on the directional cell-to-cell polar auxin transport (PAT) in various developmental contexts(*1*). Asymmetrically plasma membrane (PM)- localized auxin efflux carriers PIN-FORMED (PIN) proteins are crucial for PAT(*2, 3*) and for diverse auxin-mediated self-organization of plant patterning(*4–7*).

It has been well documented that the dynamics of PIN activity and polarity are regulated by intracellular trafficking involving clathrin-mediated PIN endocytosis(*8*) and ARF-GEF GNOM-dependent PIN endosomal recycling(*9, 10*) as well as post-translational modifications, including reversible phosphorylation(*11–13*).

Phosphorylation of PINs by various protein kinases at multiple phosphorylation sites has been implicated in the modulation of PIN polarity for the maintenance of local auxin gradients and in the regulation of PIN activity. Among PIN-phosphorylating kinases are D6 PROTEIN KINASE (D6PK)(*14*), PINOID (PID)(*15*), AGC kinases(*16*), MAP kinases(*17, 18*), CAMEL receptor-like kinase(*19*) and CPK29(*20*) which underscores the importance of PIN protein phosphorylation in plant developmental processes(*21*).

Previous work described the self-regulation of auxin transport by auxin itself as driving many plant developmental processes, such as embryo axis formation, organogenesis, vascular venation pattern and tropic responses(*22–25*). However, molecular insights into the mechanism operating between auxin-mediated PIN repolarization and PIN-based auxin transport have not been provided.

Recently, transmembrane kinases (TMKs), key components in cell surface auxin signaling, were uncovered as playing essential roles in plant development, including cell shape formation(*26*), apical hook maintenance(*27*), lateral root development(*28*), root and shoot growth(*29, 30*) and vascular tissue formation and regeneration(*31*). Notably, the *tmk1;tmk4* double mutant displayed a certain ratio of the aberrant cotyledons and fused leaf-cups, which were typically found in the *pin1-5* mutant (Fig. 1a and b), likely by affecting auxin-gradient formation during organogenesis. Further analysis showed that the embryo axis in *tmk1;tmk4* was disordered, similar to what is seen in the *pin1-5* mutant, and as compared to wild type (Figure S1). To test the possibility that TMKs are involved in the regulation of auxin transport, we used *DR5rev::GFP* auxin response reporter(*7*) in roots. Auxin treatment in roots strongly enhanced GFP signal at the root tip in the wild type, but the response was remarkably reduced in *tmk1;tmk4* (Figure S2). We further monitored the auxin distribution in the embryos of *tmk1;tmk4* as visualized by DR5 (Figure S3). The results showed that the auxin distribution at different stages of embryogenesis was disrupted in the *tmk1;tmk4* mutant compared with the wild type, which suggested TMKs are required for auxin redistribution in both roots and embryos. Then, we introduced *pPIN1::PIN1-GFP* into the *tmk1;tmk4*, and *pin1-5* mutants, respectively. PIN1 is basally localized in the stele and endodermis cells which directs auxin flow in *Arabidopsis* roots. Unlike in the wild-type roots, where PIN1-GFP was polarized at the basal side of endodermis cells, the polarity of PIN1 was significantly disrupted in the *tmk1;tmk4* double mutant with PIN1 signal at inner lateral membranes (Fig. 1 c-f), suggesting the essential role of TMKs in guiding PIN1 localization. These results show that TMKs play a vital role in both organizing the developmental patterning at the level of organs and in modulating PIN1 polarization at the level of individual cells.

**Figure 1.**
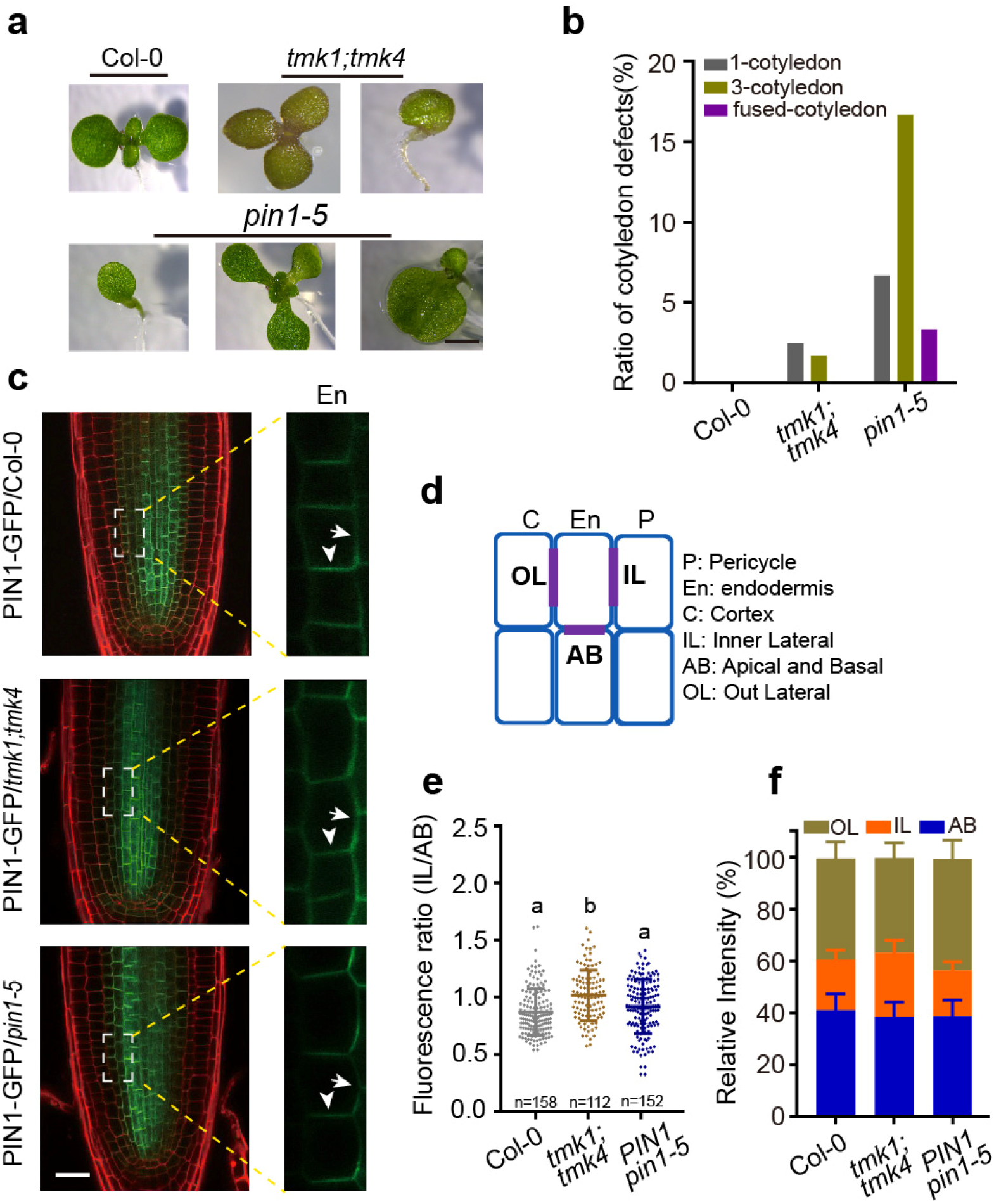
TMKs regulate PIN1 polarity and growth patterns in *Arabidopsis*. **a**, Representative figures of the cotyledons in Col-0, *tmk1;tmk4* and *pin1-5*. Scale bar, 1 mm. **b**, Quantification of the ratio of abnormal cotyledons in (***a***). **c**, Fluorescence images of PIN1-GFP in roots of Col-0, *tmk1;tmk4*, and *pin1-5*, respectively. Red signals are from PI staining and arrows indicated basally intensified and laterally depolarized localization of PIN1-GFP in endodermis (En) cells of the root. Scale bar, 25 μm. **d**, The description of the sides of root cells for quantification. **e**, Quantification of the PIN1 polarity in the endodermis shown in (***c***). The fluorescence ratio of inner lateral (IL)/apical and basal (AB) membranes sides of PIN1. **f**, The fluorescence ratio of outer lateral (OL in gray), inner lateral (IL in orange), and apical and basal (AB in blue) membrane was estimated with PIN1 in Col-0, *tmk1;tmk4* and *pin1-5* background.

To investigate the potential roles of TMK-based auxin signaling in PIN1-driven auxin transport, we introduced *pTMK1:gTMK1-RFP* into a *pPIN1:gPIN1-GFP* transgenic line. TMK1-RFP showed partial overlapping localization with PIN1-GFP at the plasma membrane of root cells (Figure S4), suggesting a direct connection between TMK1 and PIN1. By immunoprecipitating the TMK1-GFP protein from *pTMK1::TMK1-GFP* transgenic plants, we identified putative TMK1 interaction partners by mass spectrometric analysis, among which we found the PIN1 protein. Using the yeast-2-hybrid assay, we found that the hydrophilic loop of PIN1 (PIN1L) directly interacted with intracellular domains of TMK1, TMK3, and TMK4 (TMK1C, TMK3C, TMK4C) (Fig. 2a). To confirm the direct interaction, an *in vitro* pull-down assay showed that His-SUMO-TMK1C was able to bind with GST-tagged PIN1L (Fig. 2b). Moreover, Flag-tagged PIN1 was able to coimmunoprecipitate with HA-tagged TMK1 in protoplasts, and we also found that the co-IP efficiency increased with IAA treatment (Fig. 2c), suggesting an auxin-promoted TMK1-PIN1 interaction. By harboring protoplast cells in a microfluid chip(*30*), we visualized the live FRET signal between TMK1-GFP and PIN1-RFP in protoplasts after 100 nM IAA treatment. Auxin stimulated interaction between TMK1 and PIN1 within 15 seconds, suggesting a dynamic and auxin-assembled TMK1-PIN1 protein complex in live cells (Figure 2d and e).

**Figure 2.**
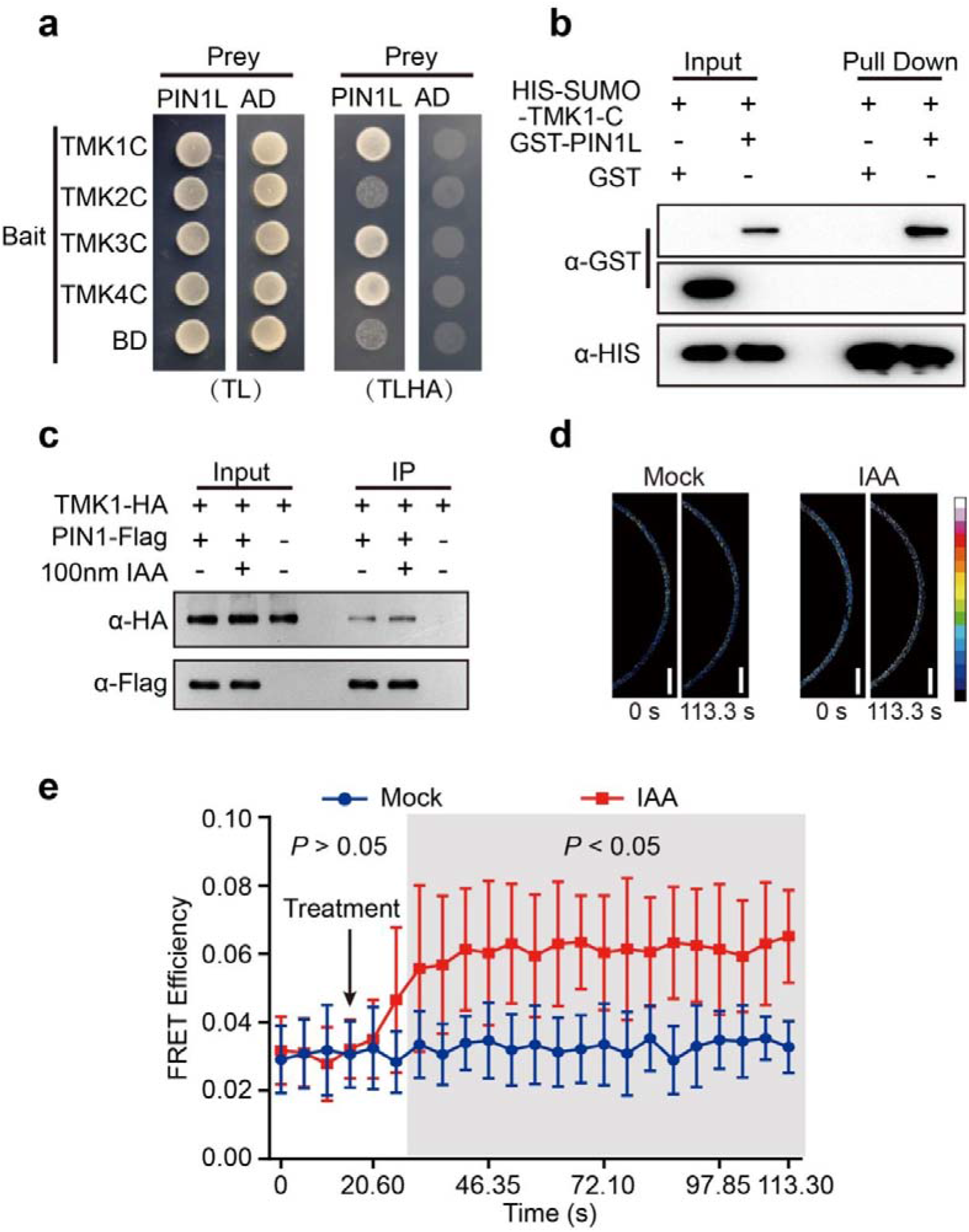
Auxin stimulates the TMK1-PIN1 complex. **a**, Y2H assay between PIN1L (hydrophilic loop) protein (as prey) and C terminus of TMKs (as bait). AD, activation domain; BD, DNA-binding domain; -TL, synthetic dextrose minimal medium without leucine and tryptophan; -TLHA, synthetic dextrose minimal medium without leucine, tryptophan, adenine, and histidine. **b**, Pull-down assay showing PIN1L-GST can be pulled down by C terminus of His-SUMO-TMK1 (TMK1-C). GST proteins were used as a negative control. TMK1-C was detected using an anti-His antibody. PIN1L-GST and GST were detected by an anti-GST antibody. **c**, TMK1 was coimmunoprecipitated with PIN1 in protoplasts. 35S-TMK1-HA and 35S-PIN1-Flag were transiently co-expressed in *Arabidopsis* protoplasts with or without IAA. 35S-TMK1-HA was used as a control. **d**, FRET analysis of the interaction between TMK1-GFP and PIN1-RFP after 100 nM auxin treatment. The times before and after treatment are indicated (0□s and 113.3□s). Scale bars, 5□μm. **e**, Quantitative time-course analyses of changes in the FRET efficiencies. IAA (100□nM) or mock buffer was applied when imaging started. The error bars indicate the SD of 9-10 cells scored. Statistical analysis was performed by two-sided Student’s t-tests (P<0.05).□

Previous works suggested PIN1 phosphorylation by different kinases, and we thus further tested whether TMK1, as a receptor-like kinase, had the ability to phosphorylate PIN1L. We performed an *in vitro* kinase assay, which incubated the His-tagged hydrophilic loop (HL) of PIN1 with His-MBP-TMK1C (C-terminus with the kinase domain) and used ^32^P-ATP as a phosphate donor. The phosphorylation reaction showed that TMK1C could phosphorylate PIN1L directly (Fig. 3a and b). To elucidate further the regulation of PIN1 by TMKs, we analyzed the phosphorylation sites in the PIN1 proteins by LC-MS/MS after incubating with the TMK1 kinase. The amino acids at S214, S252, S253, T257, S261, T449, and T458 of PIN1 were phosphorylated by TMK1 (Fig. 3a and b). Notably, three of these sites (S252, S253, and T257), we found, were shared with previously reported protein kinases PID, CPK29, and CAMEL, respectively(*19, 20, 32*). The substitution of serine and threonine by alanine at these seven sites of PIN1L (PIN1L-7A) that simulated the non-phosphorylated version of proteins led to a dramatic reduction TMK1-mediated phosphorylation *in vitro* (Fig. 3a and b), indicating that seven sites of PIN1L are targeted by TMK1. Collectively, our data demonstrate that TMK1 directly interacts with the auxin efflux carrier PIN1 and phosphorylates its hydrophilic loop.

**Figure 3.**
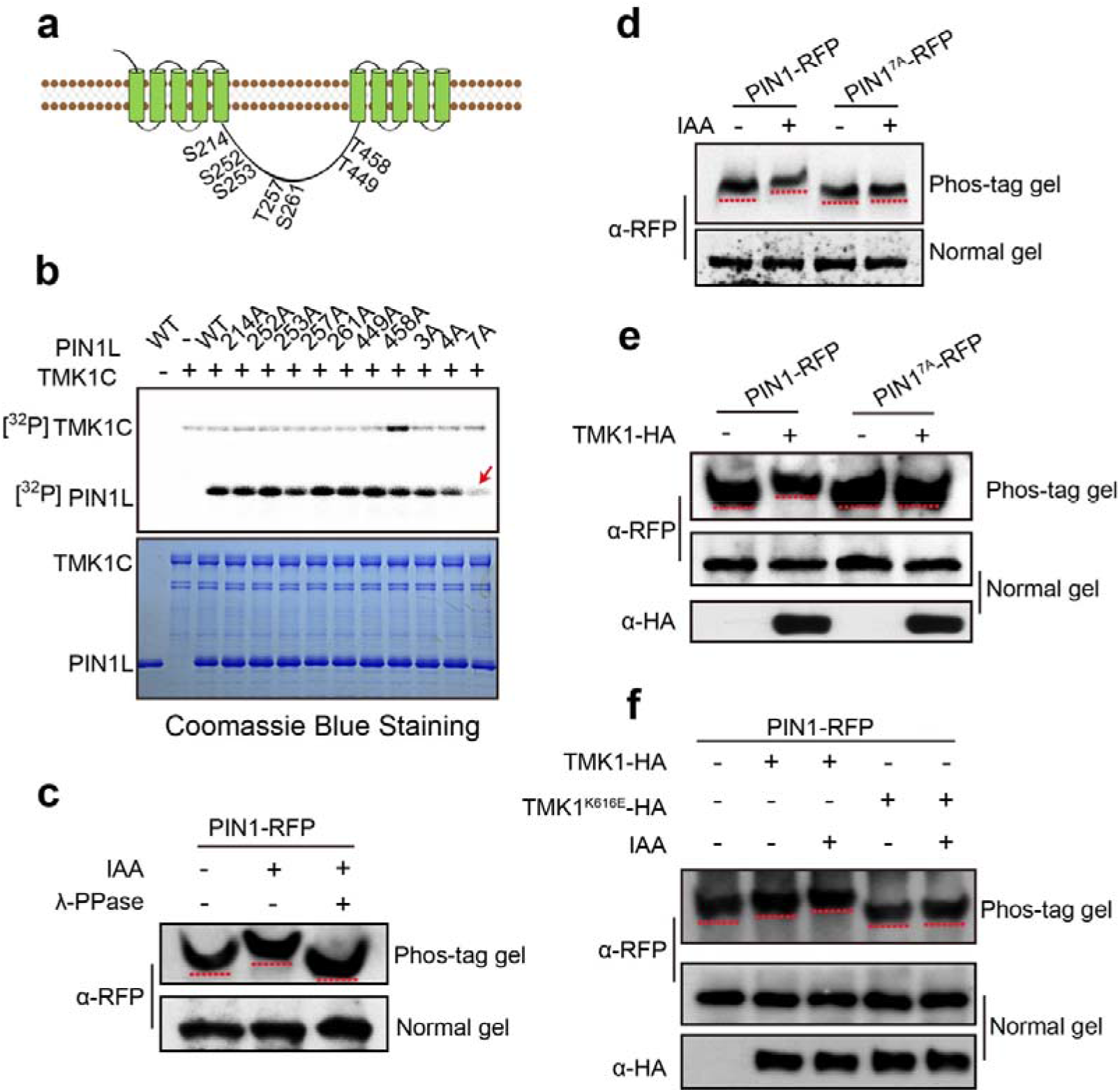
Auxin-TMK node triggers PIN1 phosphorylation. **a**, Schematic representation of PIN1 in the plasma membrane, with marked positions of TMK1-targeted phosphosites in the hydrophilic loop. **b**, *In vitro* kinase assay of the C-terminus of TMK1 with PIN1L protein. Phosphorylation is detected by radioactive P^32^. Protein loading is shown with Coomassie blue staining. 3 A: including S214A, T449A, and T458A; 4A: including S252A, S253A, T257A, and S261A; 7A: including all seven putative phosphosites. **c**, Auxin modulates the phosphorylation status of PIN1. The 35S-PIN1-RFP was transiently expressed in *Arabidopsis* protoplasts for 5 h and followed by 100 nM IAA treatment for 1 h. PIN1 protein was detected with an anti-RFP antibody. **d**, Auxin triggers PIN1 phosphorylation through TMK1-targeted sites. The 35S-PIN1-RFP and 35S-PIN1^7A^-RFP were transiently expressed in *Arabidopsis* protoplasts for 5 h and followed by 100 nM IAA treatment for 1 h, respectively. PIN1 protein was detected with an anti-RFP antibody. **e**, TMK1 induces the phosphorylation of PIN1, not PIN1^7A^. The 35S-PIN1-RFP or 35S-PIN1^7A^-GFP with 35S-TMK1-HA were transiently expressed in *Arabidopsis* protoplasts for 6 h and total protein was separated in a phos-tag SDS-PAGE. **f**, TMK1 and its kinase activity are required for the induction of PIN1 phosphorylation by auxin. The 35S-PIN1-RFP with 35S-TMK1-HA or 35S-TMK1(K616E)-HA were transiently expressed in *Arabidopsis* protoplasts for 5 h and followed by 100 nM IAA treatment for 1 h. Total protein was separated in a phos-tag SDS–PAGE and immunodetected by an anti-RFP antibody (top). The immunoblot was performed to detect PIN1 (middle) and TMK1 (bottom) in a normal SDS-PAGE as input.

Given that TMKs mediate auxin signaling at the cell surface, we further tested whether auxin regulates the PIN1 phosphorylation status in a TMKs-dependent manner. We assessed PIN1 phosphorylation through a phos-tag gel blot, in which the phosphorylated proteins specifically bind to the phos-tag reagent, causing reduced mobility compared with the non-phosphorylated counterparts. PIN1-RFP was transiently expressed in protoplasts and immunoprecipitated for the phos-tag assay after auxin treatment. Auxin was able to decrease the mobility of PIN1 proteins in phos-tag gels, which was recovered by Lambda Protein Phosphatase (λ-PPase) treatment, demonstrating that auxin induces PIN1 phosphorylation in plant cells (Fig. 3c).

To test whether auxin induces PIN1 phosphorylation via TMKs, we substituted the seven TMK1-targeted phosphorylation sites in PIN1 with alanine to simulate a dephosphorylated PIN1 (PIN1^7A^) *in vivo*. Notably, auxin was unable to promote PIN1^7A^ phosphorylation *in vivo* compared to PIN1 (Fig. 3d). This suggested that auxin regulates PIN1 phosphorylation through TMK1-targeted sites. Next, we coexpressed TMK1-HA with PIN1-RFP or PIN1^7A^-RFP in *Arabidopsis* protoplasts. TMK1 reduced the mobility shift of PIN1 in a phos-tag gel, but did not affect PIN1^7A^, suggesting that TMK1-stimulated PIN1 phosphorylation relies on the phosphorylation sites we identified (Fig. 3e). This was further supported by a kinase-dead TMK1^K616E^, a mutant with lysine 616 replaced by glutamate. TMK1^K616E^ did not promote PIN1 phosphorylation compared to TMK1 and led to a reduced auxin sensitivity of PIN1 phosphorylation (Fig. 3f).

Taken together, these results support that TMKs are responsible for auxin-promoted phosphorylation of PIN1 *in vivo*.

To further test whether phosphorylation of PIN1 by TMKs is required for PIN1 function *in planta*, we transformed phospho-dead PIN1^7A^ (mutated to alanine) and phospho-mimic PIN1^7D^ (mutated to aspartic acid) constructs under native PIN1 promoter into the *pin1-5* background. PIN1^7A^-GFP exhibited a depolarized localization at the basal side of the endodermis cells similar to PIN1-GFP in the *tmk1;tmk4* double mutant (Fig. 4a and b). PIN1^7D^ accumulated at both basal and lateral sides, which also partially disrupted PIN1 polarity (Fig. 4a and b). Consistent with its localization defects, PIN1^7A^-GFP was less able to complement the disordered cotyledon phenotype of *pin1-5* (Fig. 4c and d), while PIN1^7D^ could partially rescue the phenotype. These results again indicated the importance of PIN1 phosphorylation by TMKs in plants.

**Figure 4.**
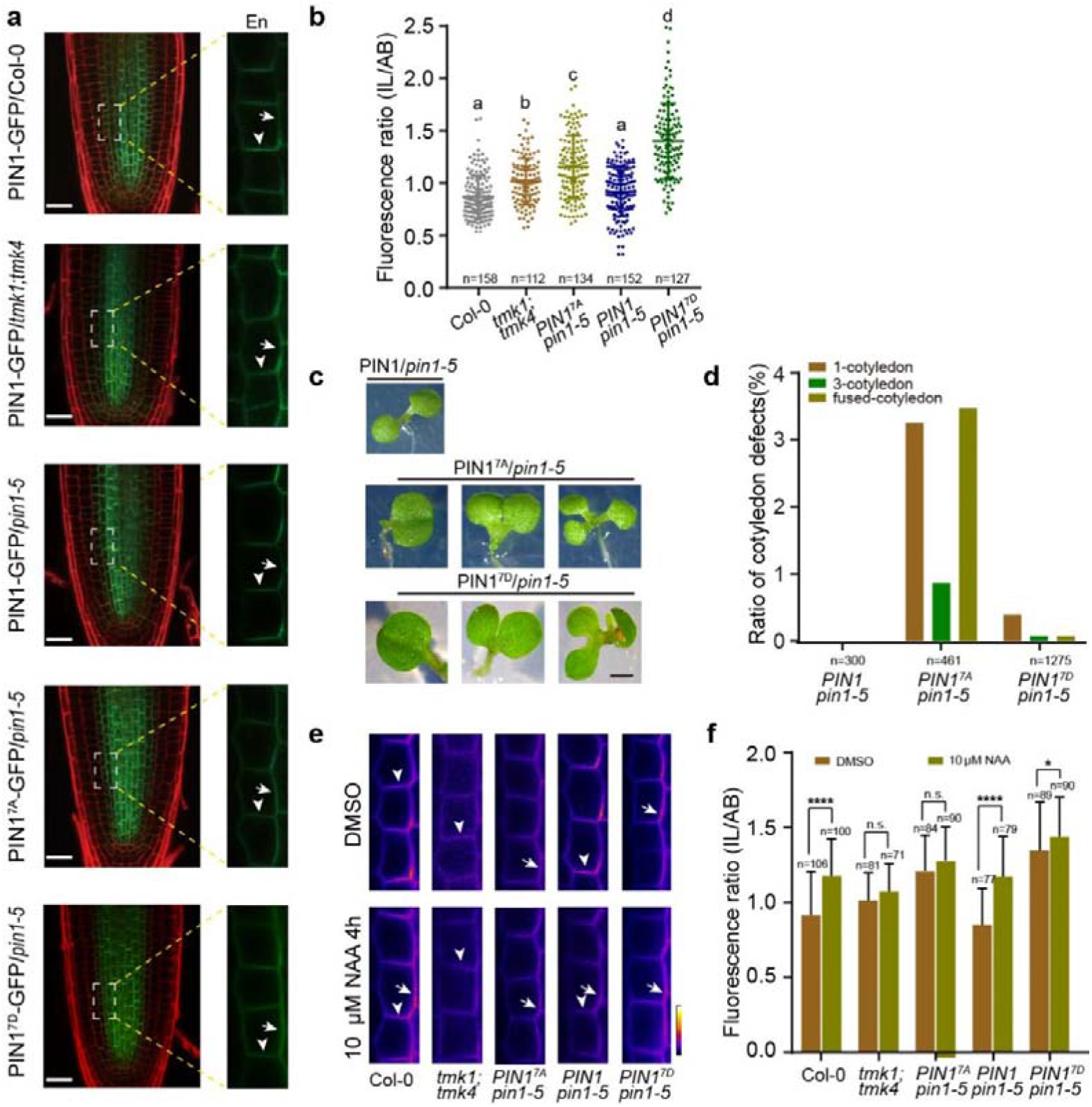
TMK-targeted phosphosites in the PIN1 are required for auxin feedback on PIN1 polarity. **a**, TMK-targeted phosphosites in the PIN1 are required for PIN1 polar localization. Representative confocal images of PIN1-GFP/Col-0, *PIN1-GFP/tmk1;tmk4*, PIN1^7A^-*GFP/pin1-5*, and PIN1^7D^-GFP/ *pin1-5* primary root, respectively. Red signals were from PI staining and arrows indicated basally intensified and laterally depolarized localization of PIN1-GFP in endodermis (En) cells of the root. Scale bar, 25 μm. **b**, Quantification of the PIN1 polarity in the endodermis shown in (a). The fluorescence ratio of inner lateral (IL)/apical and basal (AB) membranes sides. n denotes the number of scored cells. Different letters denote significant differences from one-way ANOVA with Tukey multiple comparisons test. **c**, Representative images of cotyledon phenotypes in *pPIN1::PIN1-GFP/pin1-5, pPIN1::PIN1^7A^-GFP/pin1-5* and *pPIN1::PIN1^7D^-GFP/pin1-5*, 5 days old. **d**, Quantification of the ratio of cotyledon defects shown in (a). n denotes the number of evaluated seedlings. **e**, TMK-targeted phosphosites in the PIN1 are required for the auxin feedback on PIN1 repolarization. Subcellular localization of PIN1-GFP in endodermis of root after 4 hours of 10 μM naphthaleneacetic acid (NAA) treatment. **f**, Quantitative evaluation of (e) shows the mean ratio of PIN1 IL to AB signal intensity ratio in endodermal cells. Error bars indicate the standard error of the mean. A one-way ANOVA test compared marked datasets (****P < 0.0001, n.s, no significant difference).

Auxin has been shown to be required to maintain PAT capacity and to direct itself transport by changing subcellular PIN polarity(*33*). Auxin feedback on PIN polarity relied on its constitutive endocytic recycling(*34, 35*), which promoted us to test whether TMKs are involved in auxin-induced PIN1 polarity rearrangements. Auxin was able to promote PIN1-GFP lateralization, however, this effect was partially abolished in the *tmk1;tmk4* mutant. Both PIN1^7A^-GFP and PIN1^7D^-GFP showed resistance to auxin referring to lateralization and less repolarization after auxin treatment (Fig. 4e and f). Consistently, both PIN1 -GFP/*tmk1;tmk4* and PIN1^7A^-GFP showed resistance to BFA treatment (Fig S5). All these data collectively suggest that auxin regulates PIN1 subcellular trafficking through a TMK-based mechanism, which further determines proper PIN1 localization in plant cells.

Altogether, our work uncovered an essential mechanism in regulating PIN1-driven auxin transport by a cell surface-triggered auxin signaling. This decent regulatory mechanism relies on the direct phosphorylation of PIN1 protein by TMKs, by which a short-distance auxin gradient would feedback to direct auxin flux for coordinated the polarization of individual cells throughout the tissue. Our finding provides a direct mechanism for the self-regulation of auxin transport by auxin itself, which act as a key premise for self-organization features of auxin in the regulation of plant development.

## Materials and Methods

### Plant materials and growth conditions

The Columbia 0 (Col-0) was used as a wild type. *tmk1;tmk4* double mutants, which were generated from *tmk1* (SALK_016360) crossed with *tmk4* (GABI_348E01), and *pin1-5* in Col-0 background were used for phenotypic analysis. All plant materials were grown in a plant growth chamber (PERVICAL AR-66L3) at 22 □ on half-strength Murashige and Skoog (1/2 MS) medium (pH 5.8) with 0.8% (w/v) agar and stratified in the dark at 4 °C for 3 days followed under a 16-h light/8-h dark photoperiod.

### Yeast Two-Hybrid Assay

The C-terminus of TMK family genes (*TMK1C, TMK2C, TMK3C*, and *TMK4C*) were cloned and inserted into the pGBKT7 vector as bait(*1*). The hydrophilic loop of PIN1 was amplified from *Arabidopsis* cDNA and cloned into the pGADT7 vector as prey. And the procedures have been described previously(*2*).

### Protoplast Transient Expression Assay

Protoplast transient expression assays were performed as previously described(*3*). For experiments analyzing the effect of IAA on PIN1 phosphorylation activities, protoplasts (6×10^4^) were transfected with 20 μg PIN1-RFP for 5 hours and then treated with 100 nM IAA for 1 hour. HBT-PIN1-RFP plasmids (20 μg) were co-transformed with HBT-TMK1-HA plasmids or HBT-TMK1(K616E)-HA (10 μg) into 300 μL (6×10^4^) protoplasts respectively. Protoplasts were incubated in WI buffer for 5 hours and then treated with 100 nM IAA for 1 hour. For the Co-IP assay between TMK1-HA and PIN1-Flag, the HBT-TMK1-HA plasmids (35 μg) were co-transformed with HBT-PIN1-Flag (65 μg) into 900 μL (1.8×10^5^) protoplasts, then the protoplasts were incubated in WI buffer for 6 hours.

### Co-immunoprecipitation Assay

Co-immunoprecipitation assays were performed using the *Arabidopsis* protoplast system. Protein extracted from the protoplasts transformed or co-transformed with plasmids PIN1-Flag and TMK1-HA were lysed in 500 μL of lysis buffer (50 mM Tris-HCl pH 7.4, 75 mM NaCl, 5 mM EDTA, 0.5% TritonX-100, 1×protease cocktail inhibitors). For immunoprecipitation, protein extracts were incubated with Flag Agarose beads at 4 □ for 3 hours. The immunoprecipitated proteins were washed three times with washing buffer (50 mM Tris-HCl pH 7.4, 75 mM NaCl, 5 mM EDTA) before SDS-PAGE separation and protein blot analyses, and then detected by the respective primary antibodies (Anti-Flag, Chromotek; Anti-HA, Abmart).

### Recombinant protein expression and purification

Transfected *E. coli* BL21(DE3) in 600 mL LB medium with ampicillin (100 μg/mL) were grown at 37 °C until A600 = 0.7. The cell culture was then cooled to 18 °C and recombinant protein expression was induced by IPTG adding to a final concentration of 0.3 mM at 18 °C for 12 hours. Cells were harvested by centrifugation at 4,000×g for 15 min and stored at −80 °C. Cells expressing His-SUMO tagged proteins were lysed by sonication in lysis buffer (50 mM Tris-HCl pH 7.4, 150 mM NaCl, 5 mM EDTA, and 20 mM Imidazole). The lysed solution was centrifuged at 12,000 × g for 30 min and the supernatant was incubated with Ni-NTA agarose (Qiagen) at 4 □ for 3 hours. The Ni-NTA agarose was washed three times with the wash buffer (50 mM Tris-HCl pH 7.4, 150 mM NaCl, 5 mM EDTA, and 40 mM Imidazole). The His-SUMO protein was eluted from the beads by elution buffer (50 mM Tris-HCl pH 7.4, 150 mM NaCl, and 200 mM Imidazole) and concentrated by Amicon Ultra-4 Centrifugal Filter Unit (Merck). The GST-tagged protein cells were lysed by the sonication in the equilibration buffer-phosphate-buffered saline (10 mM phosphate buffer pH 7.4, 150 mM NaCl). The lysed solution was centrifuged at 12,000 × g for 30 min and the supernatant was incubated with Glutathione agarose (Sigma) at 4 □ for 3 hours. The Glutathione agarose was washed three times with the equilibration buffer. The GST protein was eluted from the beads by elution buffer (50 mM Tris-HCl pH 9.0, 10 mM reduced glutathione (Sigma)) and concentrated by Amicon Ultra-4 Centrifugal Filter Unit (Merck).

### Pull-down assay

His-SUMO-TMK1 protein was expressed in *E.coli* BL21 and captured by Ni-NTA resin. PIN1L (166-464 aa) sequence was amplified and inserted into pGEX4T-2 vector. GST-PIN1L or GST proteins were purified from BL21 by Glutathione agarose (sigma G4510) and incubated with His-SUMO-TMK1 beads in binding buffer (20 mM HEPES, 611 pH7.5, 40 mM KCl, 1 mM EDTA, 1% glycerol, 0.1‰ TritonX-100 with PMSF) for 2 hours at 4 °C. The beads were washed three times by washing buffer (20 mM HEPES, pH7.5, 40 mM KCl, 1 mM EDTA, 0.05‰ TritonX-100). GST-PIN1L and GST proteins were detected by GST antibody (Abmart M20007), and His-SUMO-TMK1 was detected by His antibody (Abmart M30111) on western blots.

### Mobility shift assay

For phosphatase treatment, total protein was extracted from *Arabidopsis* protoplasts with 1×NEB buffer (50 mM HEPES, 100 mM NaCl, 2 mM DTT, 0.01% Brij 35) added with 1 mM MnCl_2_ and 0.5% Triton X-100. The protein extracts were incubated with Lambda Protein Phosphatase (λ-PPase) (NEB, Cat. No. P0753S) at 30 □ for 30 min and subsequently added 5×SDS sample buffer incubated at 95 □ for 5 min. Proteins were separated on a 6% SDS-PAGE gel containing 50 μM phos-tag reagent (Wako, Cat. No. 304-93525) and 100 μM ZnCl2. The separated proteins were then transferred to a nitrocellulose membrane and blotted with an anti-RFP antibody (Abcam No. 34767) at a dilution of 1/2,000.

### In vitro kinase assay

For *in vitro* kinase assays, His-MBP-TMK1C protein and mutant His-PIN1L proteins with various mutations were mixed in 50 μL kinase buffer (5 mM HEPES, 10 mM MgCl2, 10 mM MnCl2, 1 mM DTT with 2 μCi of [γ^32^P] ATP and 10 μM ATP) at 25 □ for 1-hour incubation, respectively. The reactions were stopped by adding an SDS-PAGE loading sample buffer. After separation on SDS-PAGE and gel drying, radioactive signals from His-PIN1L were detected on the dried gel by the Typhoon imaging system (GE Healthcare TYPHOON FLA 9500).

### Fluorescence imaging

PIN1-GFP and *DR5rev::GFP* signal in root and embryo were visualized in water without fixation. GFP signals were detected by a Leica SP8 confocal microscope (Leica TCS SP8 X, excitation 488 nm; emission 500-550 nm) with HyD. RFP and Propidium Iodide (PI) signals in roots were detected by a Leica SP8 confocal microscope (Leica TCS SP8 X, excitation 561 nm; emission 570-700 nm) with PMT.

### Quantitative analysis of PIN1 repolarization

Quantitative analysis of PIN1 relocalization was performed as previously described(*4*).

The mean fluorescence intensity of the PIN1 signal at the inner lateral and the basal membrane of endodermis cells was measured using Image J software. The quotients between lateral and basal values were calculated for 3-6 cells of one root and averaged. The averages of at least 15 roots per treatment were subsequently averaged again. The values of each genotype were standardized to the corresponding untreated control. Error bars in all graphs indicate standard error. Actual values are given in Supplementary Table 1.

### FRET analysis of protoplasts captured in microfluidics chips

FRET assay was performed in the *Arabidopsis* protoplasts system. TMK1 and PIN1 were fused to GFP and RFP, respectively, to generate receptor and donor proteins. Before this assay, the protoplast capture chip was filled with the WI solution (4 mM pH 5.7 MES-KOH, 0.5 M mannitol and 20 mM KCl). Using 75% ethanol to entirely remove air bubbles from the chip, then the protoplast suspension was injected slowly into the chip. The time-lapse FRET images were acquired at 5.15 s per frame using the Leica TCS SP8X confocal laser scanning microscope.

GFP and RFP signal was detected by WLL (white light laser) laser with excitation at 488 nm and 561 nm, respectively. Fluorescent emissions from 500 to 550 nm for GFP and from 570 to 640 nm for RFP were collected with HyD Gating (0.3-12 ns). When the live imaging started, IAA or mock solution was injected into the chip from the inlet. The FRET efficiency was analyzed using FRET-sensitized emission methods(*5*). A segmented line was drawn along the PM region to measure the mean signal intensity for each channel using Image-Pro Plus. The correction factors α, β, γ, and δ were calculated with the donor- and acceptor-only reference samples and the FRET efficiency was calculated as follows (*6*).

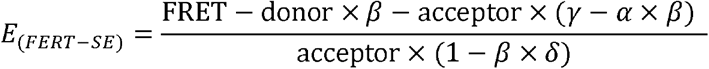

The mean FRET efficiency was calculated from 9-10 cells of 100□nM NAA or mock treatment. To generate the FRET efficiency heat-map image, the plasma membrane region captured from the donor, FRET and acceptor channels were cropped as the region of interest, and the cropped images were processed using the image calculator module of ImageJ with the *E_(FRET-SE)_* equation shown above.

### Embryo imaging

Different developmental stages of the embryo were monitored by DIC (differential interference contrast microscopy) (Nikon Ni-U) after clear solution (chloral hydrate: H_2_O: glycerol = 8:3:1) treatment for 1 hour.

### BFA-induced internalization of the PIN1 protein

5 days-old seedlings were treated with 50 μM BFA (Sigma-Aldrich; B7651) for 90 min, and the BFA-induced internalization of PIN1::GFP was observed with a confocal microscope.

### Statistical analysis

All bar graphs were generated by GraphPad Prism 7 software. Statistical significances based on *t*-test and one-way ANOVA were determined with the GraphPad Prism 7.0 (GraphPad Software, http://www.graphpad.com).

## Acknowledgment

We thank Lukáŝ Fiedler for helping with the writing. This work was supported by the National Natural Science Foundation of China (Grant 32130010, 31422008), and startup funds from both FAFU and PSC to T. X.

## Data availability

Data supporting the findings of this study are available within the paper and its Supplementary materials.

## Supplementary Materials

**Figure S1.**
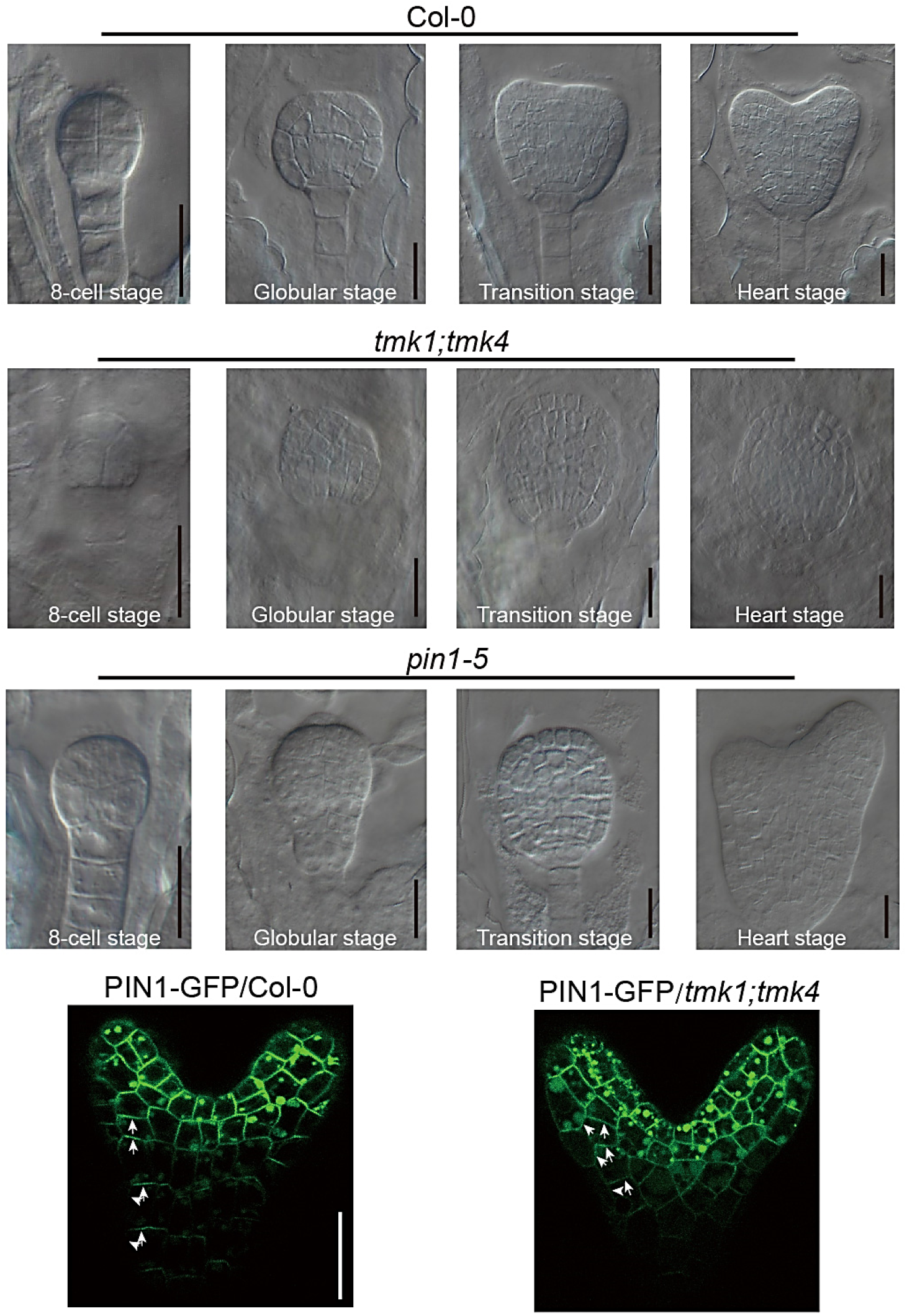
The disordered embryo axis and PIN1 mislocalization in *tmk1;tmk4*. **a,** Cell growth pattern in the embryos of Col-0, *tmk1;tmk4* and *pin1-5* at different stages. Scale bar, 20 μm. **b,** PIN1 localization in the embryos of Col-0 and *tmk1;tmk4*. Scale bar, 25 μm.

**Figure S2.**
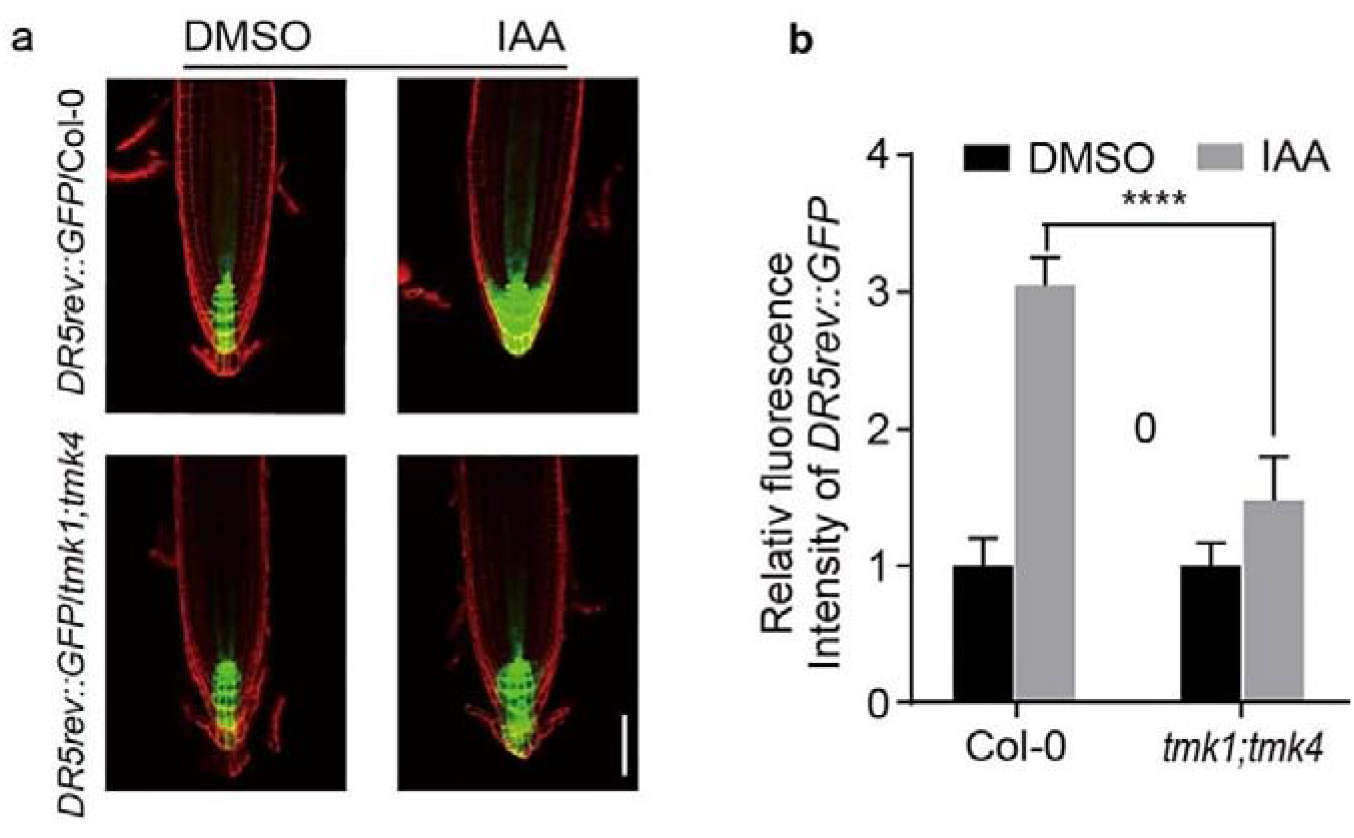
TMKs are required for auxin redistribution in the root. **a,** Expression patterns of *DR5rev::GFP* in Col-0 and *tmk1;tmk4* seedlings. The seedings *DR5rev::GFP/Col-0* and *DR5rev::GFP/tmk1;tmk4* are grown on 1/2MS for 5 days, then transferred to a medium containing 100 nM NAA for another 20 hours. The *DR5rev::GFP* shows a high auxin level in the root tip. The image was captured by Leica SP8 microscope. Scale bar, 50 μm. **b,** Quantification of relative fluorescence intensity of *DR5rev::GFP* in Col-0 and *tmk1;tmk4* background with or without NAA treatment of (A), through image J software. Error bars represent SD; Asterisks indicate a statistically significant difference (****, P value < 0.001), according to Student’s *t*-test.

**Figure S3.**
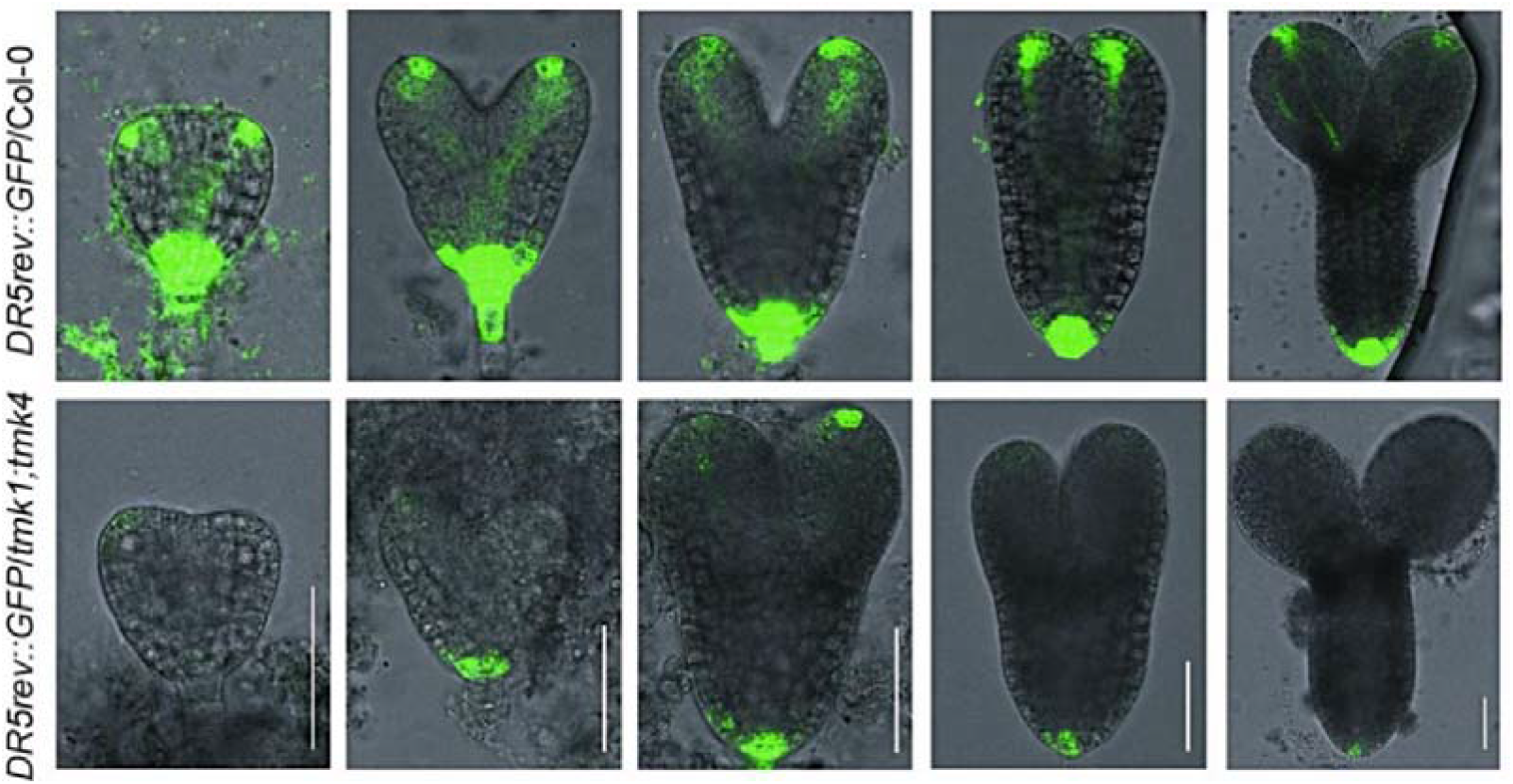
TMKs are required for auxin redistribution in the embryo. The distribution patterns of *DR5rev::GFP* in the embryos of Col-0 (up) and *tmk1;tmk4* (down) at different stages. Scale bar, 50 μm.

**Figure S4.**
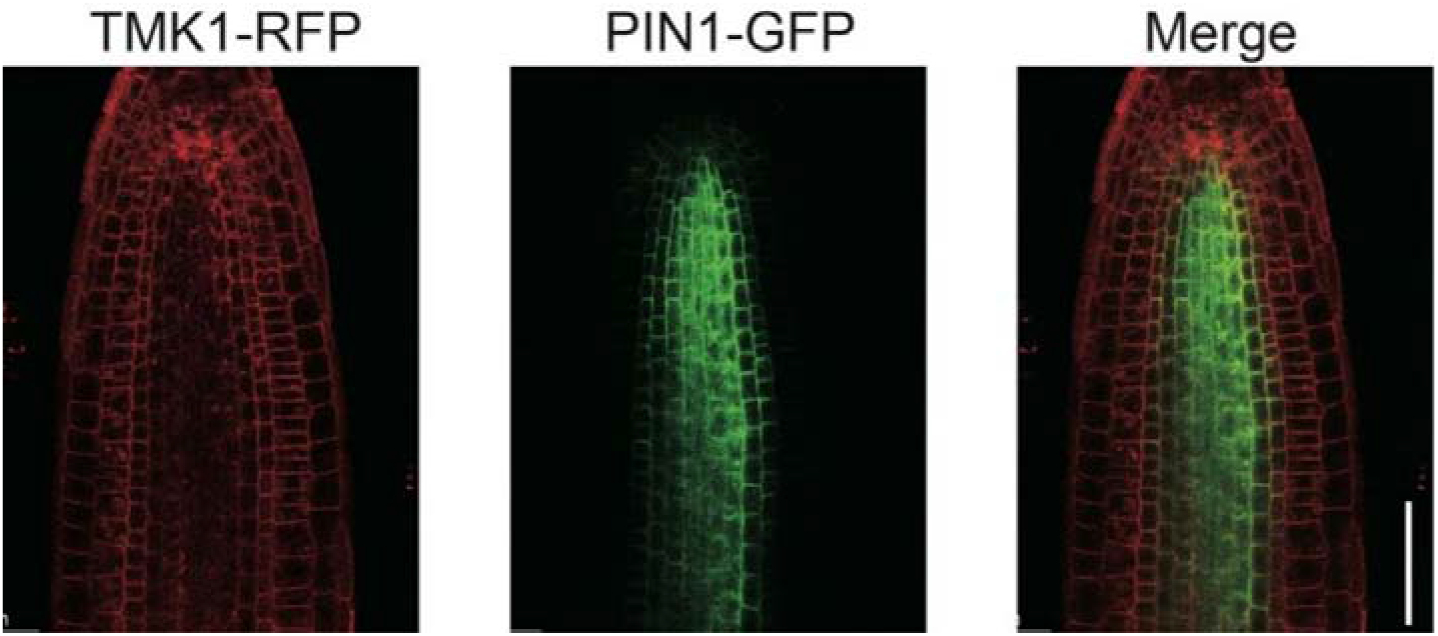
Partial co-localization between TMK1-RFP and PIN1-GFP in the root. *pTMK1::gTMK1-RFP* crossed with *pPIN1::gPIN1-GFP* was used for fluorescence observation. Scale bar, 50 μm.

**Figure S5.**
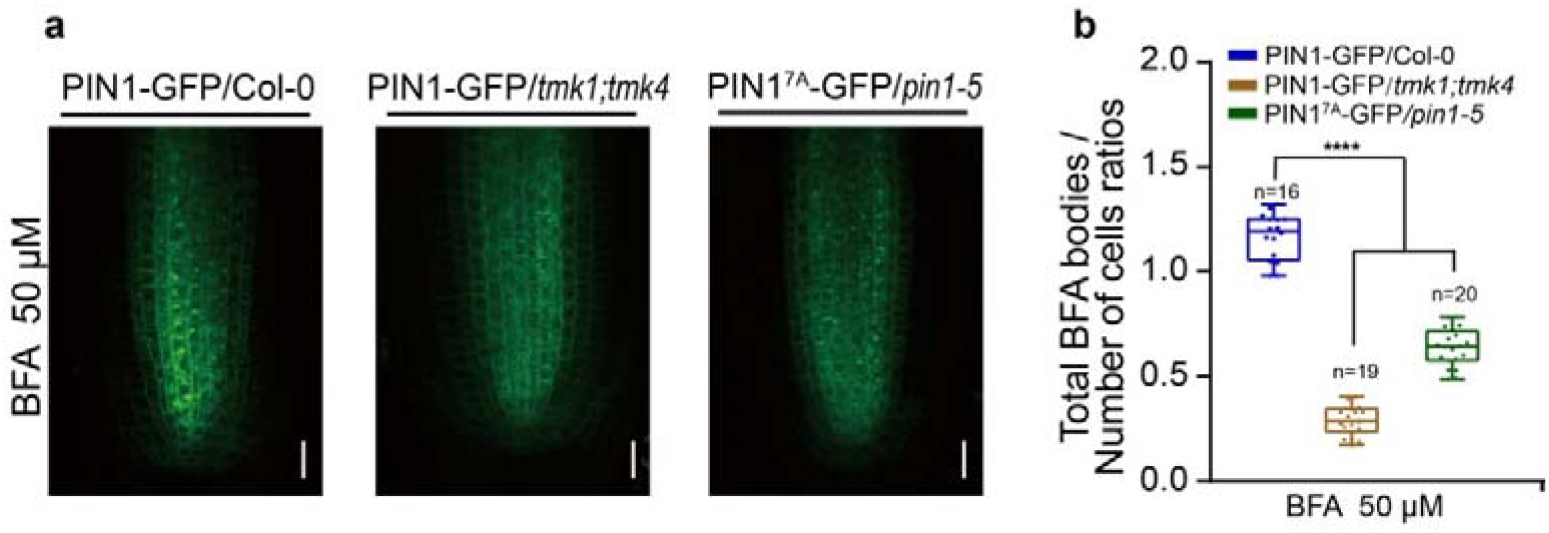
TMK-targeted phosphosites in the PIN1 are required for the PIN1 endocytic recycling. **a,** Representative confocal images of primary root stele cells after BFA-treated (50 μM) for 90 min in PIN1-GFP/Col-0, *PIN1-GFP/tmk1;tmk4* and PIN1^7A^-GFP/*pin1-5*. Scale bar, 10 mm. **b,** Quantitative evaluation of (**a**) shows the ratio of the total number of BFA bodies per total number of cells per root. n denotes the number of evaluated seedlings (**** *P* < 0.0001).

